# Multiple climate change factors jointly increase the competitiveness of a C_4_ grass against its C_3_ competitor

**DOI:** 10.1101/2022.09.01.506280

**Authors:** Gen-Chang Hsu, Po-Ju Ke, Chuan-Kai Ho

## Abstract

C_3_ and C_4_ plants are the two major terrestrial plant groups that respond differently to temperature, water, and CO_2_. Although much research has focused on the individual effect of warming, water availability, and elevated CO_2_ on biomass production and competitive interactions between C_3_ and C_4_ plants, the joint effects of these factors remain underexplored. We grew naturally co-occurring C_3_ (*Oplismenus composites*) and C_4_ (*Paspalum conjugatum*) grass under three temperature warming scenarios (control, +2°C, and +4°C), two water supply intervals (normal 2 days vs. prolonged 7 days [drought]), and two CO_2_ concentrations (ambient 400 ppm vs. elevated 800 ppm) in growth chambers and measured their above- and below-ground dry biomass in monoculture and mixture to quantify their biomass performances and competitive responses. Warming and elevated CO_2_ together enhanced the above- and/or below-ground biomass of both C_3_ and C_4_ grass. Moreover, temperature, water, and CO_2_ interacted to increase the below-ground biomass of the C_4_ grass. Surprisingly, the C_3_ grass performed worse under interspecific competition (mixture biomass) relative to intraspecific competition (monoculture biomass) at elevated CO_2_ compared to ambient CO_2_, while the reverse is true for the C_4_ grass. Furthermore, the C_4_ grass was most competitive against the C_3_ grass under simultaneous 4°C warming, drought, and elevated CO_2_. Taken together, our results suggest that the competitive balance could potentially shift in favor of C_4_ plants under the projected temperature warming, increased drought frequencies, and rising atmospheric CO_2_, with an increase in C_4_ relative abundance in future plant communities.

## Introduction

C_3_ and C_4_ plants are two major terrestrial plant groups that differ in their physiology and response to surrounding environments (Ehleringer and Monson 1993, Yamori et al. 2014). Their relative abundance in the community is closely related to various ecosystem functions and processes such as resource provisioning for herbivores, nutrient cycling, and carbon storage (Still et al. 2003, Warne et al. 2010, Pau et al. 2013). For example, C_3_ plants generally have higher nitrogen-to-carbon ratios than C_4_ plants (Barbehenn et al. 2004), and therefore their relative biomass production in the habitat may affect the nutritional availability for herbivores and even organisms at higher trophic levels (Warne et al. 2010). Importantly, because C_3_ and C_4_ plants evolved under different climates and have different adaptations to surrounding environments, understanding how they respond to climate change and the resulting dynamic shifts is crucial for the management of terrestrial ecosystems.

The performances of C_3_ and C_4_ plants under different environmental conditions can influence their relative competitive abilities (Pearcy et al. 1981, Edwards and Still 2008). Theoretical and empirical studies have suggested that C_3_ plants generally perform better and are stronger competitors in cool and well-watered environments, whereas C_4_ plants are more adapted to high temperatures and drought conditions and thus more competitive than in hot and arid environments (Teeri and Stowe 1976, Ehleringer and Monson 1993, Sage and Monson 1998, Edwards and Still 2008). In addition to temperature and water availability, C_3_ and C_4_ plants exhibit differential responses to CO_2_ (Pearcy and Ehleringer 1984, Wand et al. 1999). The photosynthesis of C_3_ plants are limited by current level of atmospheric CO_2_, and increasing CO_2_ concentration could stimulate their growth and therefore offer them competitive advantages over C_4_ plants (Ward et al. 1999, Poorter and Navas 2003, Lee 2011). These shifts in C_3_-C_4_ competitive relationships in response to climate change may alter their relative abundance, which could generate profound consequences for community and ecosystem dynamics (Ehleringer and Monson 1993, Still et al. 2003). Furthermore, different climate change variables can have counteracting effects on their competitive relationships. For example, while higher temperatures and drought conditions increase the competitive ability of C_4_ plants against C_3_ plants, elevated CO_2_ may enhance C_3_ plant performance and thus ameliorate the negative effects of warming and drought on their competitiveness, highlighting the need of simultaneously considering multiple environmental stressors for predicting C_3_-C_4_ plant dynamics under changing climate.

Different environmental factors can interact to exert a more strengthened (synergistic) or weakened (antagonistic) effect on plants than the simple summation of their individual effects (Wand et al. 1999). For example, an experiment showed a synergistic effect between water and CO_2_ treatment in C_3_-C_4_ mixture, where elevated CO_2_ had a greater positive effect on the C_4_ biomass under drought compared to well-water conditions (Valerio et al. 2011). On the other hand, another study found that the growth stimulation of elevated CO_2_ on a C_3_ crop was dampened at higher temperatures, indicating an antagonistic effect between warming and CO_2_ (Alberto et al. 1996). While extensive research has focused on the individual effect of warming, drought, and elevated CO_2_ as well as two-factor interactions on C_3_ and C_4_ plant performances and competition, studies examining all three factors simultaneously remain scarce. Because of the concurrent changes in global temperature, precipitation patterns, and atmospheric CO_2_ concentration, investigating how these three factors jointly affect C_3_-C_4_ plant dynamics could help better predict future plant communities in the face of climate change.

To examine the joint effects of the aforementioned three climate change factors on the biomass performances and competitive responses of C_3_ and C_4_ plants, we grew naturally co-occurring C_3_ (*Oplismenus composites*) and C_4_ (*Paspalum conjugatum*) grass in growth chambers under three temperature warming scenarios, two water supply intervals, and two

CO_2_ concentrations. We compared the above- and below-ground dry biomass in monoculture and mixture to assess their competitive responses under different experimental treatments. Based on the the ecophysiology of C_3_ and C_4_ plants, we predict a greater competitiveness of the C_3_ grass under lower temperatures, ample water supply, and elevated CO_2_, whereas the C_4_ grass will be more competitive under warming, drought, and ambient CO_2_.

## Materials and Methods

### Preparation of plant individuals

Ramets of *O. composites* and *P. conjugatum* were collected from the campus of National Taiwan University (N25° 00′ 57”, E121° 32′ 19”) in October, 2017. These ramets were planted in seedling nursery pots using fully-mixed peat moss soil (Kekkila natural peat moss soil) as the substrate. All plant individuals were first cultivated in greenhouse at ambient CO_2_ concentration with 25/20°C day/night temperature for two weeks, allowing their recovery and establishment prior to the start of experiment.

### Experimental design

To examine the effects of major climate change factors on the performances and competitive interactions of the C_3_ and C_4_ grass, we used a full-factorial design crossing three temperature warming scenarios, two water supply intervals, and two CO_2_ concentrations (see the following paragraph for more details). Each treatment combination contained four replicate pots of C_3_ monoculture, C_4_ monoculture, and C_3_-C_4_ mixture (each with four individuals; see the following text), yielding a total of 144 pots (12 treatment combinations × 3 planting schemes × 4 replicate pots (Fig. 1). To avoid the potential confounding effect of initial size, similar-sized individuals of each two grasses (C_3_ grass: 8.6 ± 4.3 cm; C_4_ grass: 6.2 ± 4.5 cm; mean ± SD) were selected from the greenhouse nursery for the experimental pots. Four individuals were grown and equally-spaced in each pot (13 cm in height x 12 cm in diameter), conforming to their densities at the site where the ramets were collected (personal observations). In the monoculture pots, all four individuals belonged to the same species; in the mixture pots, two individuals of each species were planted. Each replicate pot was assigned a pot number prior to the start of the experiment; monoculture and mixture pots were paired for analysis of competitive responses (Fig. 1). Pots were placed in growth chambers receiving respective treatments and a constant relative humidity of 70%, photoperiod of 12:12 light/dark, and photosynthetic photon flux density (PPFD) of 160–200 μmol · M^-2^ · S^-1^.

**Figure 1.**
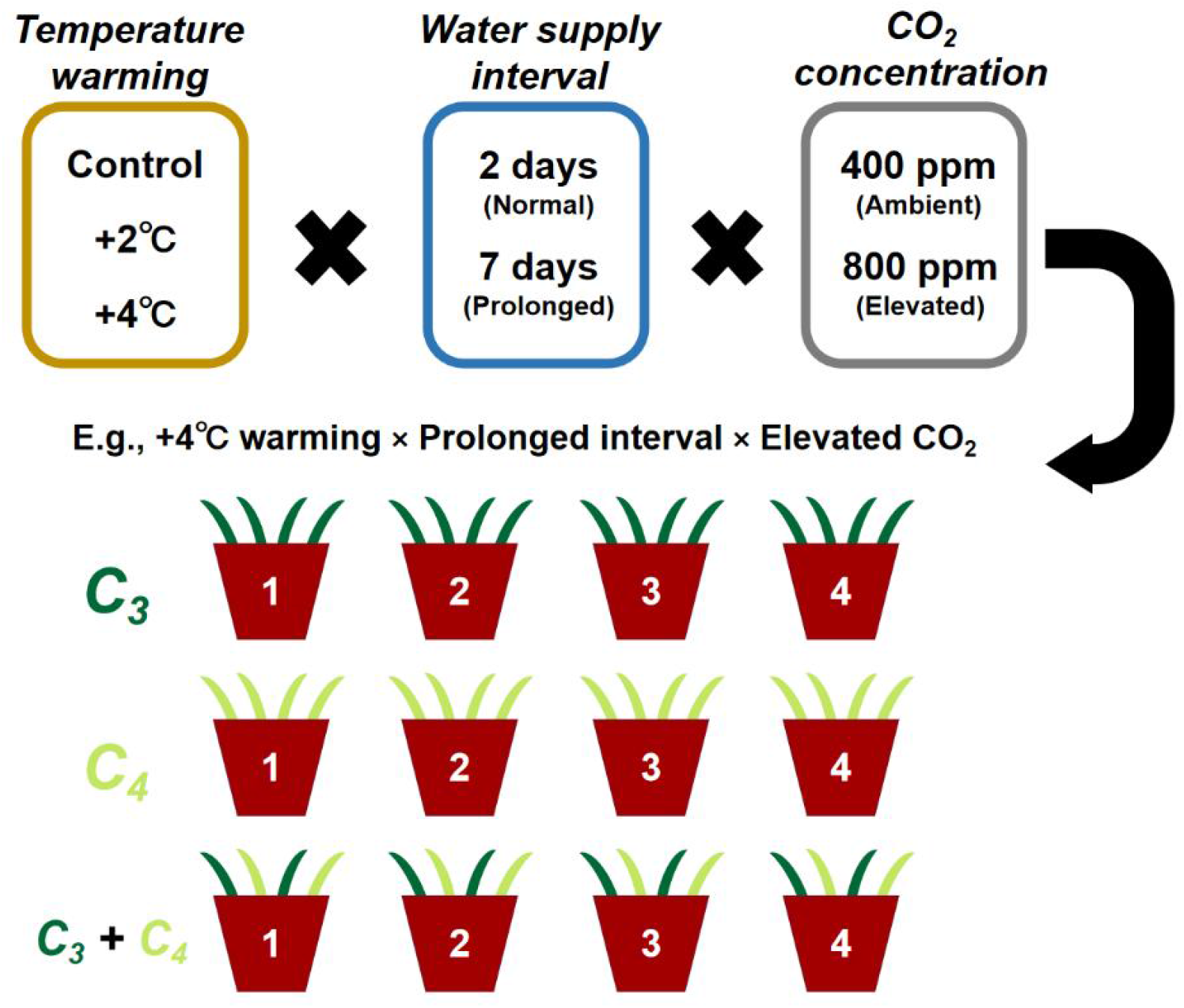
An illustration of the experimental design of this study. The experiment was conducted using a factorial design crossing three temperature warming scenarios (control, +2°C, and +4°C), two water supply intervals (normal 2 days vs. prolonged 7 days), and two CO_2_ concentrations (ambient 400 ppm vs. elevated 800 ppm). Each treatment combination contained four replicate pots of C_3_ (*Oplismenus composites*) and C_4_ grass (*Paspalum conjugatum*) in monoculture as well as C_3_-C_4_ mixture. Four plant individuals (four C_3_ or C_4_ in monoculture and two C_3_ + C_4_ in mixture) were grown in each pot. Replicate monoculture and mixture pots in each treatment combination were paired *a priori* by assigning the same pot number for analysis of competitive responses.

Three warming scenarios were used in the temperature treatment: control (+0°C), 2°C warming (+2°C), and 4°C warming (+4°C), based on the projection that mean global surface temperature will be 1.8 ± 0.5°C and 3.7 ± 0.7°C above the 1986–2005 reference period by 2100 under intermediate (Representative Concentration Pathway [RCP] 4.5) and high carbon emissions scenario (RCP 8.5), respectively (IPCC 2014). To reflect the daily temperature fluctuations in the field, a control temperature curve was derived using hourly temperature records in 2016 from the nearest weather station (Taipei weather station; mean: 23.3°C; range: 21.6–25.9°C). Temperature curves for the +2°C and +4°C treatment were obtained by horizontally shifting the control temperature curve upward by 2°C and 4°C, respectively (Fig. S1). Two watering regimes were used in the water treatment: 2 days and 7 days, representing the normal and prolonged precipitation interval (drought). The normal interval of 2 days was determined based on the mean annual number of precipitation days between 1981 and 2011 (165.5 days per year) using weather records from the Taipei weather station. The water volume for a single watering event was 160 ml (derived by multiplying the average precipitation on each rainy day [14.5 mm; mean annual precipitation of 2405 mm divided by 165.5 days] with the pot surface area [110 cm^2^]). Two CO_2_ concentrations were used in the CO_2_ treatment: 400 ppm (ambient) and 800 ppm (elevated), reflecting the current level of atmospheric CO_2_ and the projected atmospheric CO_2_ between RCP 6.0 (670 ppm) and RCP 8.5 (936 ppm) (IPCC 2014).

Plants were harvested after six weeks of cultivation and separated into above- and below-ground parts. The below-ground parts were carefully washed to remove the soil, with as many fine roots retained as possible. All plant materials were oven-dried at 50°C for seven days. Dry biomass was measured to the third decimal place in the unit of gram (g).

### Data analyses

To examine the effects of our experimental treatments on the above- and below-ground dry biomass of the C_3_ and C_4_ grass, we fitted linear mixed effects models (LMMs) with temperature warming, water supply interval, CO_2_ concentration, culture type (monoculture vs. mixture), and their interactions as fixed effects, and pot as a random effect, using the “lmer” function in the *lme4* R package (Bates et al. 2015). Models were fitted separately for the above- and below-ground biomass of the two species (i.e., a total of four models). The response (dry biomass of individual plant) was log-transformed; initial plant height was also included as a covariate. The significance of predictors was assessed via *F*-tests (type III sum of squares) using the “joint_tests” function in the *emmeans* R package (Lenth et al. 2018). Tukey post-hoc comparisons of treatment levels were conducted using also the *emmeans* R package.

To quantify the competitive responses of the C_3_ and C_4_ grass in each treatment combination, we first averaged the above-/below-ground dry biomass of the four individuals in each monoculture pot as well as the biomass of the two individuals of the same species in each mixture pot. We then calculated the log response ratio (lnRR) of the pot-level mean biomass in paired monoculture and mixture pots as log(mixture mean biomass / monoculture mean biomass). This lnRR measures the relative effects of interspecific versus intraspecific competition received by the focal individual, effectively representing the relative “sensitivity” of a species to interspecific and intraspecific competition (Bennett et al. 2016). A positive lnRR indicates a better performance under interspecific competition compared to intraspecific competition (i.e., less suppression by interspecific competition) and vice versa for a negative lnRR. The lnRRs were analyzed using linear models with temperature warming, water supply interval, CO_2_ concentration, plant species, and their interactions as predictors. The significance of predictors was assessed via *F*-tests (type III sum of squares) as above; the estimated marginal means (EMMs) of lnRRs of the two grass species derived from the linear models were compared within each treatment combination using the *emmeans* R package. All analyses were performed in R version 4.1.2 (R Core Team 2021).

## Results

### Biomass performances of the C_3_ and C_4_ grass

#### (1) C_3_ above- and below-ground dry biomass

The above-ground dry biomass of the C_3_ grass (*O. composites*) was affected by temperature warming, water supply interval, and CO_2_ concentration (Table 1). The above-ground biomass was higher under 4°C warming for all but normal water supply interval × ambient CO_2_ treatment (Fig. 2), indicating an interaction between temperature and water as well as between temperature and CO_2_ (Table 1). Prolonged water supply interval reduced the biomass, whereas elevated CO_2_ increased the above-ground biomass (Table S1).

**Table 1.** Results of the linear mixed models (LMMs) examining the effects of temperature warming, water supply interval, CO_2_ concentration, and culture type (monoculture vs. mixture) on the above-ground dry biomass of the C_3_ and C_4_ grass

| Predictor | C<sub>3</sub> grass |  |  | C<sub>4</sub> grass |  |  |
| --- | --- | --- | --- | --- | --- | --- |
|  | *F* | *df* | *P* * | *F* | *df* | *P* * |
| Temperature warming | 57.42 | 2, 103 | < **0.001** | 56.66 | 2, 99 | < **0.001** |
| Water supply interval | 26.13 | 1, 99 | < **0.001** | 1.60 | 1, 99 | 0.208 |
| CO<sub>2</sub> concentration | 48.7 | 1, 99 | < **0.001** | 18.8 | 1, 100 | < **0.001** |
| Culture type | 3.54 | 1, 99 | 0.063 | 2.30 | 1, 99 | 0.132 |
| Temperature × Water | 4.41 | 2, 99 | **0.015** | 2.82 | 2, 99 | 0.064 |
| Temperature × CO<sub>2</sub> | 18.07 | 2, 101 | < **0.001** | 3.10 | 2, 99 | **0.049** |
| Temperature × Culture type | 1.07 | 2, 99 | 0.346 | 4.89 | 2, 100 | **0.009** |
| Water × CO<sub>2</sub> | 5.74 | 1, 99 | **0.018** | 0.73 | 1, 99 | 0.394 |
| Water × Culture type | 0.39 | 1, 99 | 0.534 | 0.25 | 1, 99 | 0.619 |
| CO<sub>2</sub> × Culture type | 0.26 | 1, 102 | 0.613 | 4.57 | 1, 99 | **0.035** |
| Temperature × Water × CO<sub>2</sub> | 0.13 | 2, 99 | 0.881 | 1.84 | 2, 99 | 0.164 |
| Temperature × Water × Culture type | 1.97 | 2, 99 | 0.145 | 2.43 | 2, 100 | 0.093 |
| Temperature × CO<sub>2</sub> × Culture type | 0.90 | 2, 100 | 0.409 | 0.15 | 2, 99 | 0.861 |
| Water × CO<sub>2</sub> × Culture type | 3.08 | 1, 99 | 0.082 | 0.96 | 1, 99 | 0.331 |
| Temperature × Water × CO<sub>2</sub> × Culture type | 4.07 | 2, 99 | **0.020** | 0.26 | 2, 99 | 0.769 |

\* Significant effects ( $P < 0.05$ ) are highlighted in bold.

**Figure 2.**
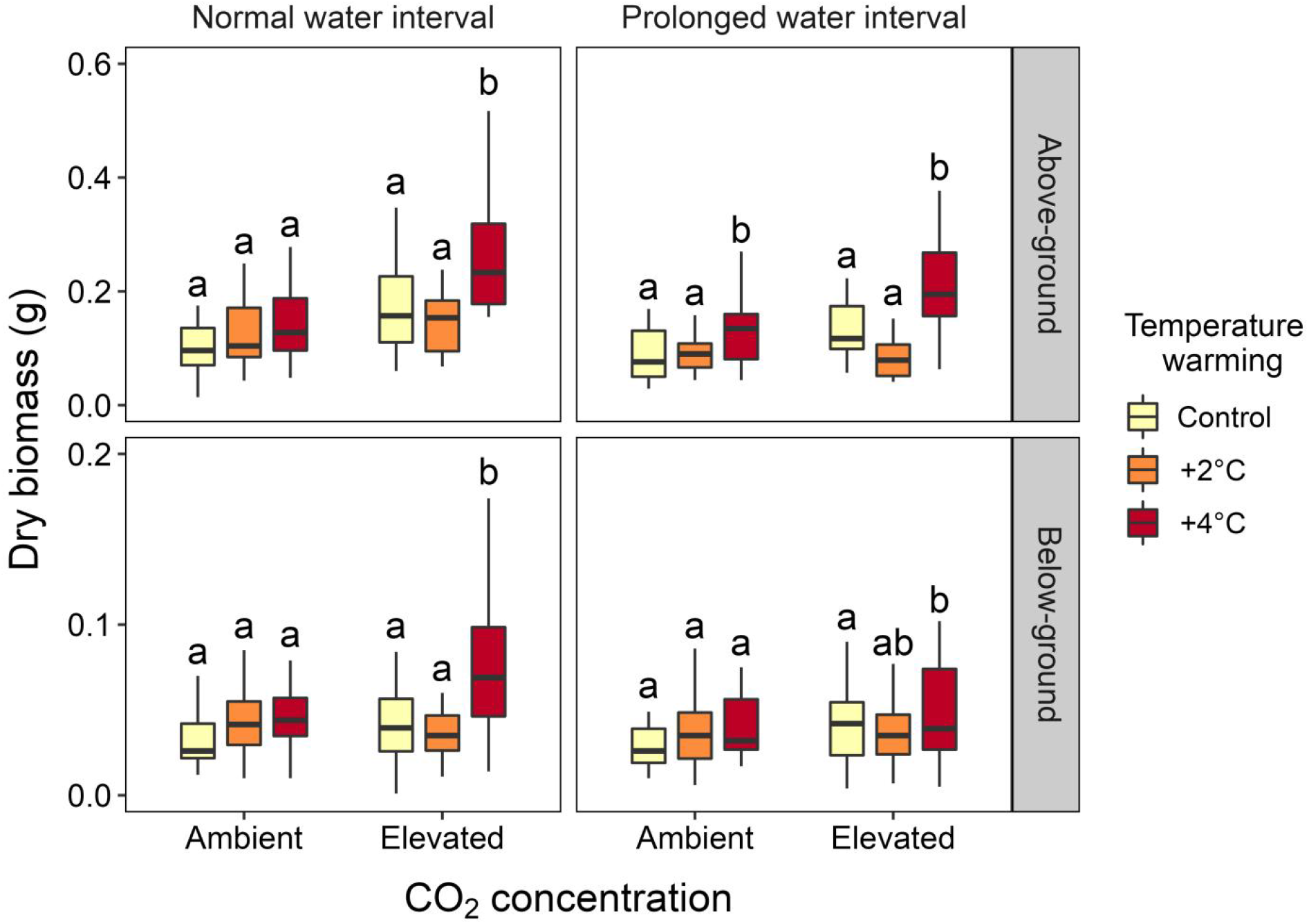
Above- and below-ground dry biomass of the C_3_ grass under different temperature warming, water supply interval, and CO_2_ concentration treatments. Tukey post-hoc comparisons were conducted for the three temperature scenarios within each water × CO_2_ treatment combination using the estimated marginal means (EMMs) from the LMMs. Different letters indicate statistical significance (*P* < 0.05).

The below-ground dry biomass of *O. composites* was affected by temperature warming and water supply interval but not CO_2_ concentration (Table S2). The below-ground biomass did not differ among temperature treatments at ambient CO_2_ but was higher under 4°C warming at elevated CO_2_ (Fig. 2), indicating an interaction between temperature and CO_2_ (Table S2). Prolonged water supply interval reduced the below-ground biomass (Table S1).

#### (2) C_4_ above- and below-ground dry biomass

The above-ground dry biomass of the C_4_ grass (*P. conjugatum*) was affected by temperature warming and CO_2_ concentration but not water supply interval (Table 1). The above-ground biomass was higher under 4°C warming across all water × CO_2_ treatments (Fig. 3). Elevated CO_2_ enhanced the above-ground biomass production (Table S1).

**Figure 3.**
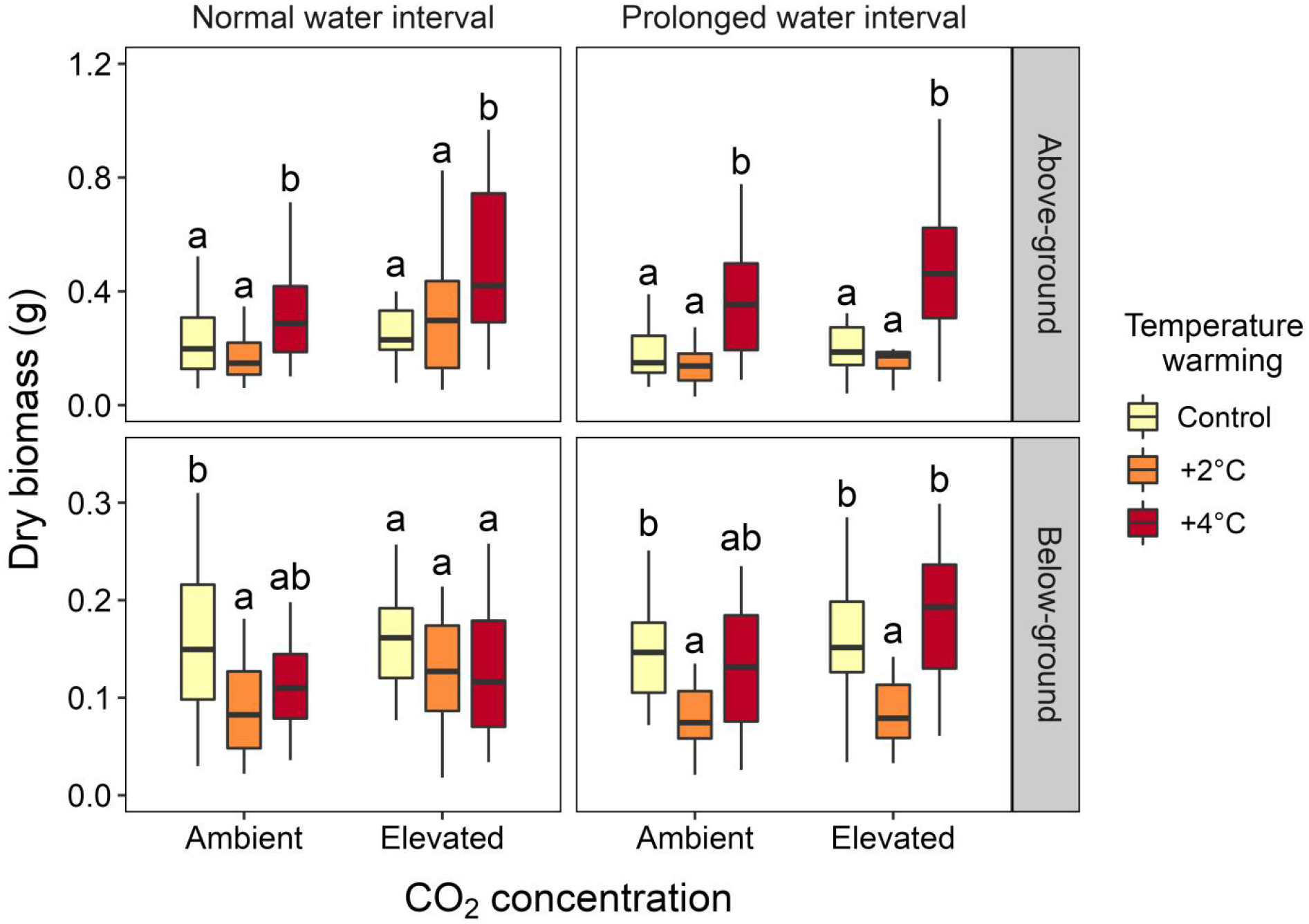
Above- and below-ground dry biomass of the C_4_ grass under different temperature warming, water supply interval, and CO_2_ concentration treatments. Tukey post-hoc comparisons were conducted for the three temperature scenarios within each water × CO_2_ treatment combination using the estimated marginal means (EMMs) from the LMMs. Different letters indicate statistical significance (*P* < 0.05).

Similar to the response of above-ground dry biomass to the experimental treatments, the below-ground dry biomass of *P. conjugatum* was affected by temperature warming and CO_2_ concentration but not water supply interval (Table S2). The effect of temperature warming on below-ground biomass varied across water and CO_2_ treatments (i.e., Temperature × Water × CO_2_ interaction; Fig. 3; Table S2). Specifically, at normal water supply interval, the below-ground biomass was lower under 2°C warming compared to control temperature at ambient CO_2_, while there was no difference in biomass among temperature treatments at elevated CO_2_. At prolonged water supply interval, the below-ground biomass was the lowest under 2°C warming, although the difference between 2°C and 4°C warming was not statistically significant at ambient CO_2_. Elevated CO_2_ enhanced the below-ground biomass production (Table S1).

When examining the effect of culture type, we found a significant Temperature × Culture type interaction for the C_4_ grass (Table 1; Table S2). In particular, the C_4_ above-ground biomass was considerably higher in mixture than in monoculture (Fig. S2), suggesting that warming may increase the competitive ability of C_4_ grass when grown with C_3_ grass.

### Competitive responses of the C_3_ and C_4_ grass

The lnRRs of above- and below-ground dry biomass were not affected by temperature warming, water supply interval, or CO_2_ concentration; however, the competitive responses of the two grasses depended on the CO_2_ concentration (Species × CO_2_ interaction; Table 2; Fig. 4). At ambient CO_2_, the C_3_ grass had similar above-ground biomass under interspecific (mixture) and intraspecific (monoculture) competition (lnRR ≈ 0) and higher below-ground biomass under interspecific competition compared to intraspecific competition (lnRR > 0), whereas the C_4_ grass had lower above- and below-ground biomass under interspecific competition compared to intraspecific competition (lnRR < 0). In contrast, the competitive responses reversed at elevated CO_2_, with the C_3_ grass having lower biomass (lnRR < 0) and the C_4_ grass having higher biomass when grown in mixtures (lnRR > 0) (Fig. 4). These results suggest that increasing CO_2_ concentration could benefit C_4_ grass when in competition with C_3_ grass.

**Table 2.** Results of the linear models examining the effects of temperature warming, water supply interval, CO_2_ concentration, and species on the log response ratios (lnRRs) of above- and below-ground dry biomass in mixture relative to monoculture (log[mixture biomass/monoculture biomass])

| Predictor | Above-ground |  |  | Below-ground |  |  |
| --- | --- | --- | --- | --- | --- | --- |
|  | *F* | *df* | *P* * | *F* | *df* | *P* * |
| Temperature warming | 0.91 | 2, 72 | 0.406 | 0.59 | 2, 71 | 0.556 |
| Water supply interval | 2.94 | 1, 72 | 0.091 | 0.40 | 1, 71 | 0.528 |
| CO<sub>2</sub> concentration | 0.22 | 1, 72 | 0.637 | 0.25 | 1, 71 | 0.621 |
| Species | 1.59 | 1, 72 | 0.211 | 0.12 | 1, 71 | 0.730 |
| Temperature × Water | 1.77 | 2, 72 | 0.177 | 1.29 | 2, 71 | 0.281 |
| Temperature × CO<sub>2</sub> | 1.36 | 2, 72 | 0.262 | 1.56 | 2, 71 | 0.218 |
| Temperature × Species | 1.98 | 2, 72 | 0.146 | 1.61 | 2, 71 | 0.207 |
| Water × CO<sub>2</sub> | 0.74 | 1, 72 | 0.392 | 2.67 | 1, 71 | 0.106 |
| Water × Species | 0.07 | 1, 72 | 0.792 | 4.97 | 1, 71 | **0.029** |
| CO<sub>2</sub> × Species | 5.66 | 1, 72 | **0.020** | 7.87 | 1, 71 | **0.006** |
| Temperature × Water × CO<sub>2</sub> | 2.87 | 2, 72 | 0.063 | 2.98 | 2, 71 | 0.057 |
| Temperature × Water × Species | 0.44 | 2, 72 | 0.645 | 0.46 | 2, 71 | 0.631 |
| Temperature × CO<sub>2</sub> × Species | 0.31 | 2, 72 | 0.736 | 0.58 | 2, 71 | 0.561 |
| Water × CO<sub>2</sub> × Species | 1.72 | 1, 72 | 0.194 | 1.43 | 1, 71 | 0.235 |
| Temperature × Water × CO<sub>2</sub> × Species | 1.60 | 2, 72 | 0.210 | 3.38 | 2, 71 | **0.039** |

\* Significant effects ( $P < 0.05$ ) are highlighted in bold.

**Figure 4.**
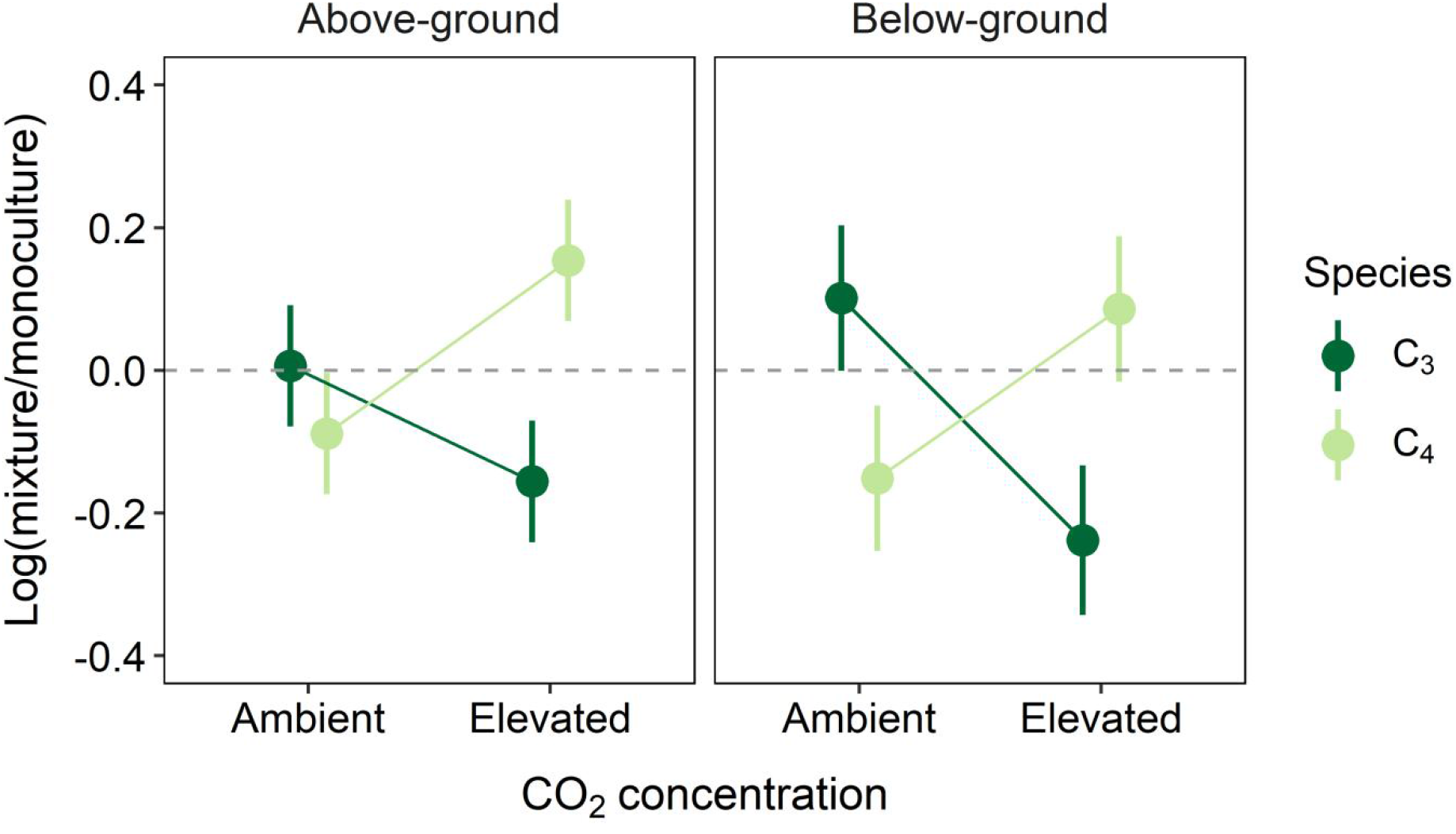
Interactions between CO_2_ concentration and species for the lnRRs of above- and below-ground dry biomass. Points and error bars represent the EMMs and standard errors derived from the linear models. Grey dashed lines (lnRR = 0) indicate equal effects of interspecific and intraspecific competition; a positive lnRR indicates a better performance under interspecific competition compared to intraspecific competition (i.e., weaker suppression by interspecific competition) and vice versa for a negative lnRR.

Overall, the above- and below-ground competitive responses of the two grass species in each treatment combination were generally comparable, despite some variations in their signs and magnitudes (Fig. S3). However, the lnRRs of both above- and below-ground dry biomass did differ between the two species under a combination of 4°C warming, prolonged water supply interval, and elevated CO_2_, where the C_3_ grass performed worse (lnRR < 0) and the C_4_ grass performed better (lnRR > 0) in mixtures compared to their corresponding monocultures (Fig. 5a; Fig. S3). Indeed, the positive lnRRs of the C_4_ grass under multiple environmental stressors were largely due to an increase in mixture biomass rather than a reduction in monoculture biomass in comparison to the controlled baseline treatment (Fig. 5b). Together, these results indicate a greater competitive ability of the C_4_ grass under higher temperature, drought condition, and elevated CO_2_.

**Figure 5.**
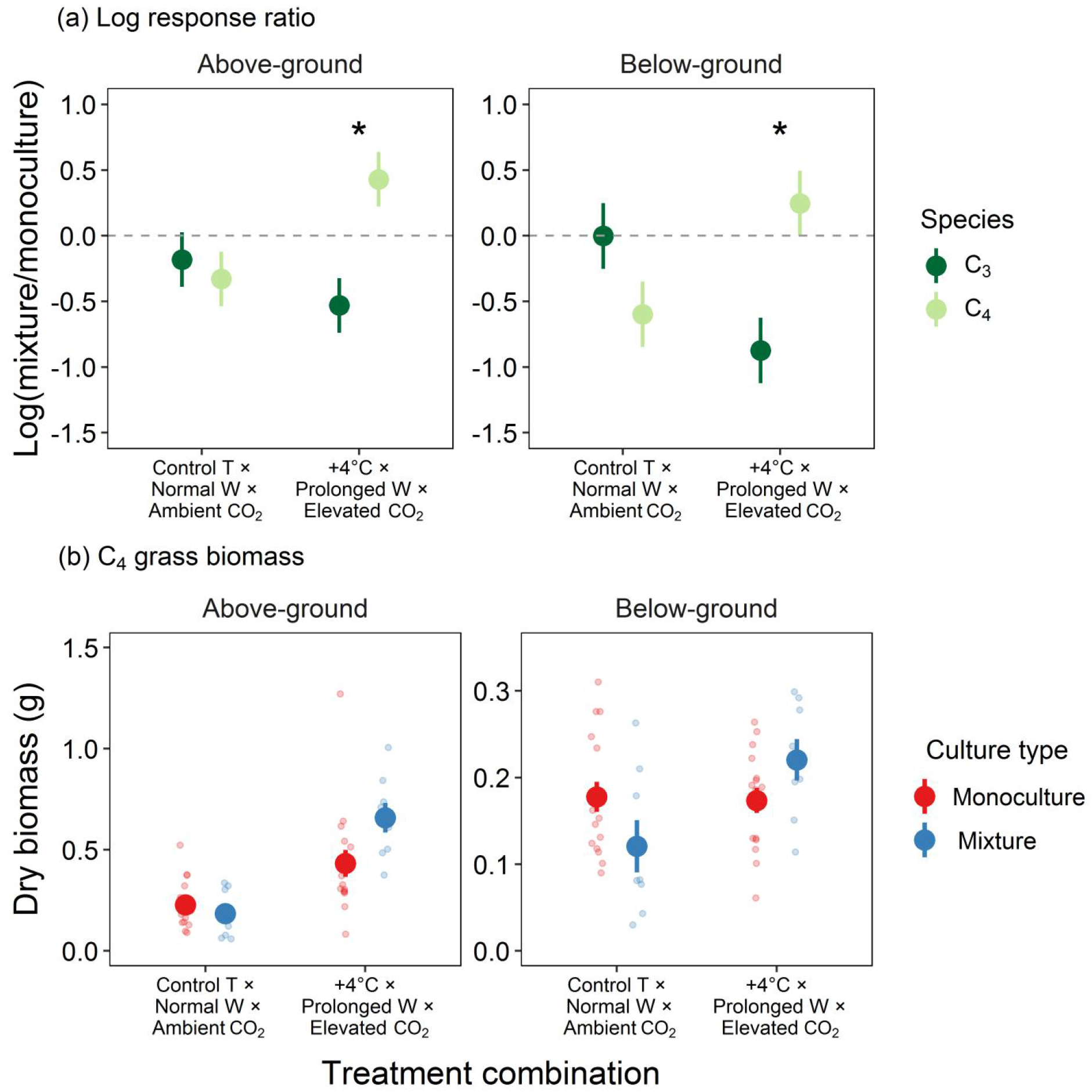
LnRRs of above- and below-ground dry biomass of the two grass species (a) and above- and below-ground dry biomass of the C_4_ grass (b) under two opposite treatment combinations: (1) control temperature × normal water supply interval × ambient CO_2_, and (2) 4°C warming × prolonged water supply interval × elevated CO_2_. Points and error bars in (a) represent the EMMs and standard errors derived from the linear models; points and error bars in (b) represent original means and standard errors. Asterisks denote a significant difference in lnRRs between the two grass species (*P* < 0.05).

## Discussion

We examined the joint effects of three major climate change factors—temperature, water, and CO_2_—on the biomass performances and competitive responses of a C_3_ and a C_4_ grass in growth chambers. Warming and elevated CO_2_ generally enhanced the above- and/or below-ground biomass of both grasses, with a synergism between these two factors. In contrast to theoretical predictions and experimental results (Carter and Peterson 1983, Pearcy and Ehleringer 1984), the C_3_ grass performed worse under interspecific competition (mixture biomass) relative to intraspecific competition (monoculture biomass) at elevated CO_2_, while the reverse is true for C_4_ grass. Furthermore, the C_4_ grass outperformed the C_3_ grass most under simultaneous 4°C warming, drought conditions, and elevated CO_2_, suggesting that the competitive balance could potentially shift in favor of C_4_ grass under the projected temperature warming, increased drought frequencies, and rising atmospheric CO_2_. Taken together, our results provide empirical evidence for C_3_ and C_4_ biomass performances and competition in response to multiple environment stressors, underlining the importance of considering multi-variable effects and offering useful insights into C_3_-C_4_ plant dynamics under future climate change.

### Effects of warming and elevated CO_2_ on biomass performances

We found that warming (in particular 4°C warming) increased the above- and below-ground biomass of the C_3_ grass, contradicting the common notion that C_3_ grass performance should not be enhanced, or even potentially be reduced, under higher temperatures (Sage and Kubien 2007). A possible explanation for this observation is that the control temperature used in the study was below the thermal optimum of photosynthesis for the C_3_ grass and therefore warming indeed led to a further increase in carbon fixation and biomass production. Additionally, the C_3_ grass may shift its thermal optimum as the temperature rises, as research has suggested that C_3_ plants have broader thermal ranges and higher potential for thermal acclimation (Sage and Kubien 2007, Yamori et al. 2014). Consequently, future climate warming might not necessarily suppress C_3_ plants as generally predicted if they are able to exhibit plastic thermal acclimation to higher temperatures.

Elevated CO_2_ enhanced the above-ground biomass of the C_3_ grass as well as the above- and below-ground biomass of the C_4_ grass. The higher C_3_ grass biomass at elevated CO_2_ agrees with the widely-documented positive response of C_3_ plants to increasing CO_2_ (Ainsworth and Long 2005, Wang et al. 2012). In contrast, biomass enhancement of the C_4_ grass is surprising as theory predicts that increasing CO_2_ concentration may not benefit the performance of C_4_ plants because of their lower CO_2_ saturation points for photosynthesis (Sage 1994, Ghannoum et al. 2000). Nonetheless, empirical evidence regarding the effect of CO_2_ on the performance of C_4_ plants is mixed. In fact, some studies have reported biomass stimulation by CO_2_ in C_4_ plants (e.g., Owensby et al. 1993, Wand et al. 1999). These findings indicate that the response of C_4_ plants to CO_2_ could be species-specific and a future rise in atmospheric CO_2_ may potentially benefit certain C_4_ plants.

### Interactions among treatment factors

The interaction between temperature and CO_2_ appeared to be the most conspicuous two-way interaction among the three treatment factors across both grass species (Table 1; Table S2). The synergistic effect of temperature warming and elevated CO_2_ suggests that climate warming together with increasing atmospheric CO_2_ could further enhance the biomass production of plants regardless of their photosynthetic pathways (Wang et al. 2012). For the C_3_ grass, such a synergism may arise from a greater ameliorating effect of CO_2_ on photorespiration with rising temperature (Long 1991, Dusenge et al. 2019). Furthermore, elevated CO_2_ could increase the thermal optimum of net carbon assimilation rate of the C_3_ grass (Sage and Kubien 2007), leading to greater biomass production under simultaneous warming and elevated CO_2_. On the other hand, C_4_ plants generally do not respond strongly to rising CO_2_ because of their CO_2_ concentration mechanism. The potential mechanisms underlying the observed positive temperature × CO_2_ interaction in the C_4_ grass warrant further investigations.

Our results revealed a three-way interaction among temperature, water, and CO_2_ for the below-ground biomass of the C_4_ grass. There are a few possible explanations for this. First, warming-induced physiological changes (e.g., reduced stomatal conductance) could provide cross tolerance to drought (Havaux 1992). Second, elevated CO_2_ may increase water use efficiency and thus help alleviate the impact of water stress (Xu et al. 2013). Third, more resources may be allocated to below-ground growth to increase water uptake in response to temperature warming and water limitation (Malik et al. 1979, Sharp and Davies 1989). Research on how C_3_ and C_4_ plants respond to multiple stressors has been limited, and our finding provides experimental evidence that future climate change could affect plant biomass production in a complex way that might not be predicted from individual factors alone. More studies on how multiple climate change factors jointly affect C_3_ and C_4_ plant performances are needed to verify if such a three-way interaction is a common phenomenon.

### Competitive responses under experimental treatments

Countering previous studies showing greater competitive advantages of C_3_ plants at elevated CO_2_ (Marks and Strain 1989, Ziska 2001), our results show that higher CO_2_ concentration could potentially benefit the C_4_ grass when in competition with the C_3_ grass. Specifically, the C_3_ grass was suppressed more by interspecific competition relative to intraspecific competition at elevated CO_2_ compared to ambient CO_2_, whereas the C_4_ grass performed better under interspecific competition relative to intraspecific competition at elevated CO_2_ compared to ambient CO_2_ (Fig. 4). Such patterns can be partly explained by a greater effect of elevated CO_2_ on the C_4_ above-ground biomass in mixture compared to monoculture (CO_2_ × Culture type interaction; Table 1). Moreover, elevated CO_2_ combined with 4°C warming and drought resulted in the largest difference in the competitive responses of the two grasses, with an increased interspecific competitive ability of the C_4_ grass against the C_3_ grass (Fig. 5). These results suggest that the competitive balance could potentially shift in favor of C_4_ plants under future climate scenarios, which may have potential impacts on both natural and agricultural systems. For example, C_4_ plants are generally less nutritious than C_3_ plants, and increasing C_4_ relative abundance in the habitat could reduce the nutritional availability for herbivores (Barbehenn et al. 2004). Furthermore, C_3_ crops may suffer greater yield loss because of increasing competition pressure from C_4_ weeds (Korres et al. 2016).

Despite some variations in the signs and magnitudes, the competitive responses of the two grass species were by and large similar in most treatment combinations, indicating similar relative effects of interspecific versus intraspecific competition received by the two grasses. Surprisingly, we did not find any main effect of temperature, water, or CO_2_ treatment on the above- and below-ground competitive responses of the two grasses, despite strong effects of these factors on their biomass production. A lack of treatment effect on the competitive responses may occur if the two grasses respond to the treatments in a similar fashion (e.g., positive responses of both C_3_ and C_4_ grass to elevated CO_2_), and consequently there might be no competitive shift because neither species would gain a net advantage over the other. This shows that performances of individual species alone may not necessarily reflect their competitive relationships, which depend largely on the species’ relative performances when grown together.

### Conclusions

Using an experimental approach, our study shows that warming and elevated CO_2_ together enhanced the biomass performances of both C_3_ and C_4_ grass, and temperature, water, and CO_2_ interacted to affect the C_4_ biomass. Moreover, simultaneous warming, drought, and particularly elevated CO_2_, benefited the C_4_ grass most when in competition with the C_3_ grass, suggesting potential shifts in their competitive balance along with an increase in C_4_ relative abundance under future climate scenarios. Changes in C_3_-C_4_ community composition can have profound consequences for the functioning of natural and agricultural systems, including primary productivity, nutrient cycling, herbivore nutritional balance, and crop production. As the effects of climate change are multifaceted and often involve complex interactions and feedbacks over time and space, long-term and large-scale field experiments involving more species will represent a promising research avenue that scales up our laboratory results to better predict C_3_-C_4_ plant dynamics under changing climate.

## Acknowledgements

We thanked William J-A Ou and Jia-Zhen Lin for assisting with experimental work and data collection. This study was supported by funding from Ministry of Science and Technology, Taiwan (106-2813-C-002-163-B).

