## Appendix S1 for "Multiple climate change factors jointly increase the competitiveness of a C_4_ grass against its C_3_ competitor"

Table S1. Contrasts between levels of water supply interval (normal 2 days − prolonged 7 days), $CO_2$ concentration (ambient 400 ppm − elevated 800 ppm), and culture type (monoculture − mixture) for above- and below-ground dry biomass of the $C_3$ and $C_4$ grass

| Species part | Treatment | Contrast† | SE | *df* | *t* | *P** |
|---|---|---|---|---|---|---|
| $C_3$ above-ground | Water | 0.23 | 0.05 | 99 | 5.11 | **< 0.001** |
| | $CO_2$ | -0.31 | 0.05 | 99 | -6.98 | **< 0.001** |
| | Culture type | 0.08 | 0.05 | 99 | 1.88 | 0.063 |
| $C_3$ below-ground | Water | 0.17 | 0.07 | 100 | 2.35 | **0.021** |
| | $CO_2$ | 0.01 | 0.07 | 100 | 0.20 | 0.844 |
| | Culture type | 0.06 | 0.07 | 100 | 0.82 | 0.415 |
| $C_4$ above-ground | Water | 0.08 | 0.07 | 99 | 1.27 | 0.208 |
| | $CO_2$ | -0.29 | 0.07 | 100 | -4.34 | **< 0.001** |
| | Culture type | -0.10 | 0.07 | 99 | -1.52 | 0.132 |
| $C_4$ below-ground | Water | -0.09 | 0.06 | 99 | -1.49 | 0.140 |
| | $CO_2$ | -0.19 | 0.06 | 100 | -2.99 | **0.004** |
| | Culture type | -0.02 | 0.06 | 99 | -0.39 | 0.697 |

† Contrasts were computed using the estimated marginal means (EMMs) from the LMMs.

* Significant effects ($P < 0.05$) are highlighted in bold.

Table S2. Results of the linear mixed models (LMMs) examining the effects of temperature warming, water supply interval, $CO_2$ concentration, and culture type (monoculture vs. mixture) on the below-ground dry biomass of the $C_3$ and $C_4$ grass

| Predictor | $C_3$ grass | | | $C_4$ grass | | |
|---|---|---|---|---|---|---|
| | *F* | *df* | *P** | *F* | *df* | *P** |
| Temperature warming | 15.87 | 2, 104 | **< 0.001** | 19.8 | 2, 99 | **< 0.001** |
| Water supply interval | 5.50 | 1, 100 | **0.021** | 2.21 | 1, 99 | 0.140 |
| $CO_2$ concentration | 0.04 | 1, 100 | 0.844 | 8.93 | 1, 100 | **0.004** |
| Culture type | 0.67 | 1, 100 | 0.415 | 0.15 | 1, 99 | 0.697 |
| Temperature × Water | 0.60 | 2, 100 | 0.552 | 3.53 | 2, 99 | **0.033** |
| Temperature × $CO_2$ | 6.34 | 2, 103 | **0.003** | 1.28 | 2, 99 | 0.281 |
| Temperature × Culture type | 0.07 | 2, 100 | 0.936 | 4.02 | 2, 100 | **0.021** |
| Water × $CO_2$ | 1.26 | 1, 100 | 0.264 | 0.10 | 1, 99 | 0.746 |
| Water × Culture type | 2.22 | 1, 100 | 0.139 | 2.91 | 1, 99 | 0.091 |
| $CO_2$ × Culture type | 1.14 | 1, 102 | 0.289 | 2.89 | 1, 99 | 0.092 |
| Temperature × Water × $CO_2$ | 1.26 | 2, 100 | 0.288 | 4.00 | 2, 99 | **0.021** |
| Temperature × Water × Culture type | 0.84 | 2, 100 | 0.433 | 1.03 | 2, 100 | 0.361 |
| Temperature × $CO_2$ × Culture type | 0.06 | 2, 101 | 0.943 | 0.56 | 2, 99 | 0.572 |
| Water × $CO_2$ × Culture type | 5.35 | 1, 100 | **0.023** | 1.41 | 1, 99 | 0.238 |
| Temperature × Water × $CO_2$ × Culture type | 3.10 | 2, 100 | **0.049** | 0.46 | 2, 99 | 0.632 |

* Significant effects ($P < 0.05$) are highlighted in bold.

Figure S1. The daily temperature curves for the three temperature warming scenarios (control, +2°C, and +4°C). See Materials and Methods for details of the curve derivation.

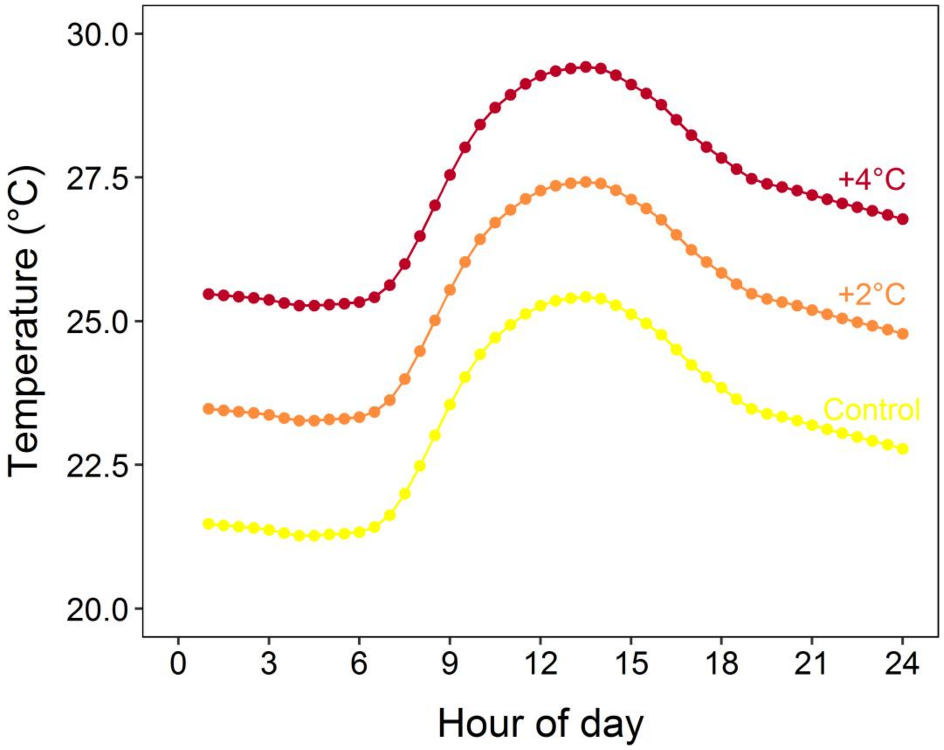

Figure S2. Above- and below-ground dry biomass of the $C_4$ grass in monoculture and mixture under different temperature warming treatments. Points and error bars represent the estimated marginal means (EMMs) and standard errors derived from the linear mixed models.

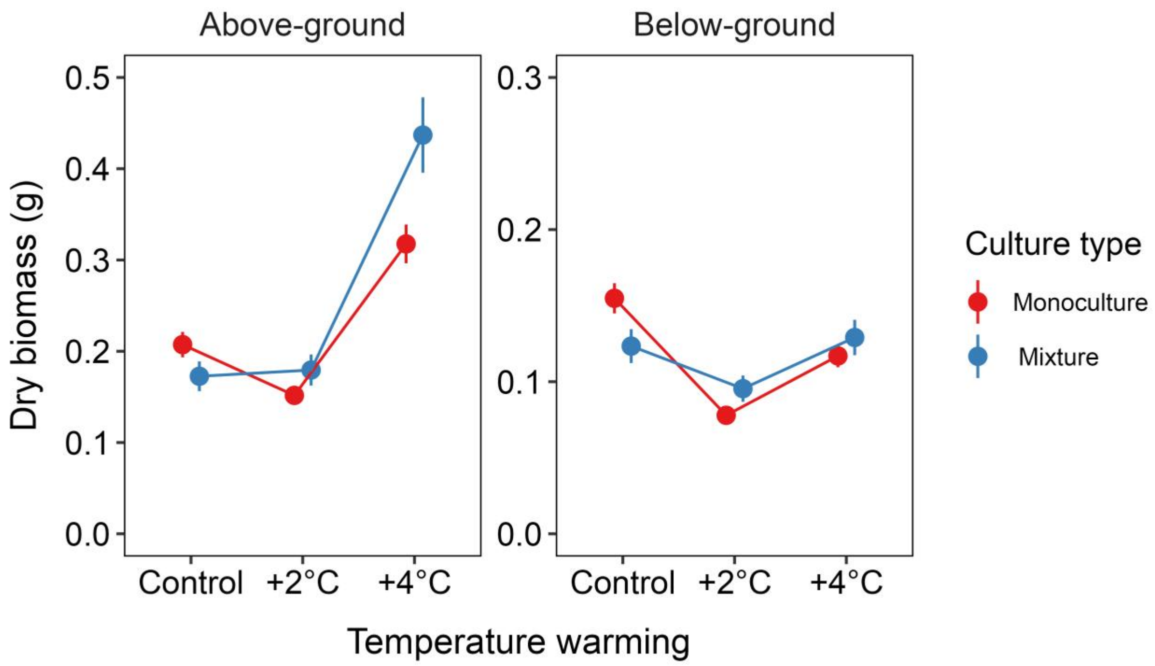

Figure S3. The log response ratios (lnRRs) of (a) above-ground and (b) below-ground dry biomass for the $C_3$ grass and the $C_4$ grass under different temperature warming, water supply interval, and $CO_2$ concentration treatments. Points and error bars represent the estimated marginal means (EMMs) and standard errors derived from the linear models. Asterisks denote a significant difference in lnRRs between the two grass species ($P < 0.05$).

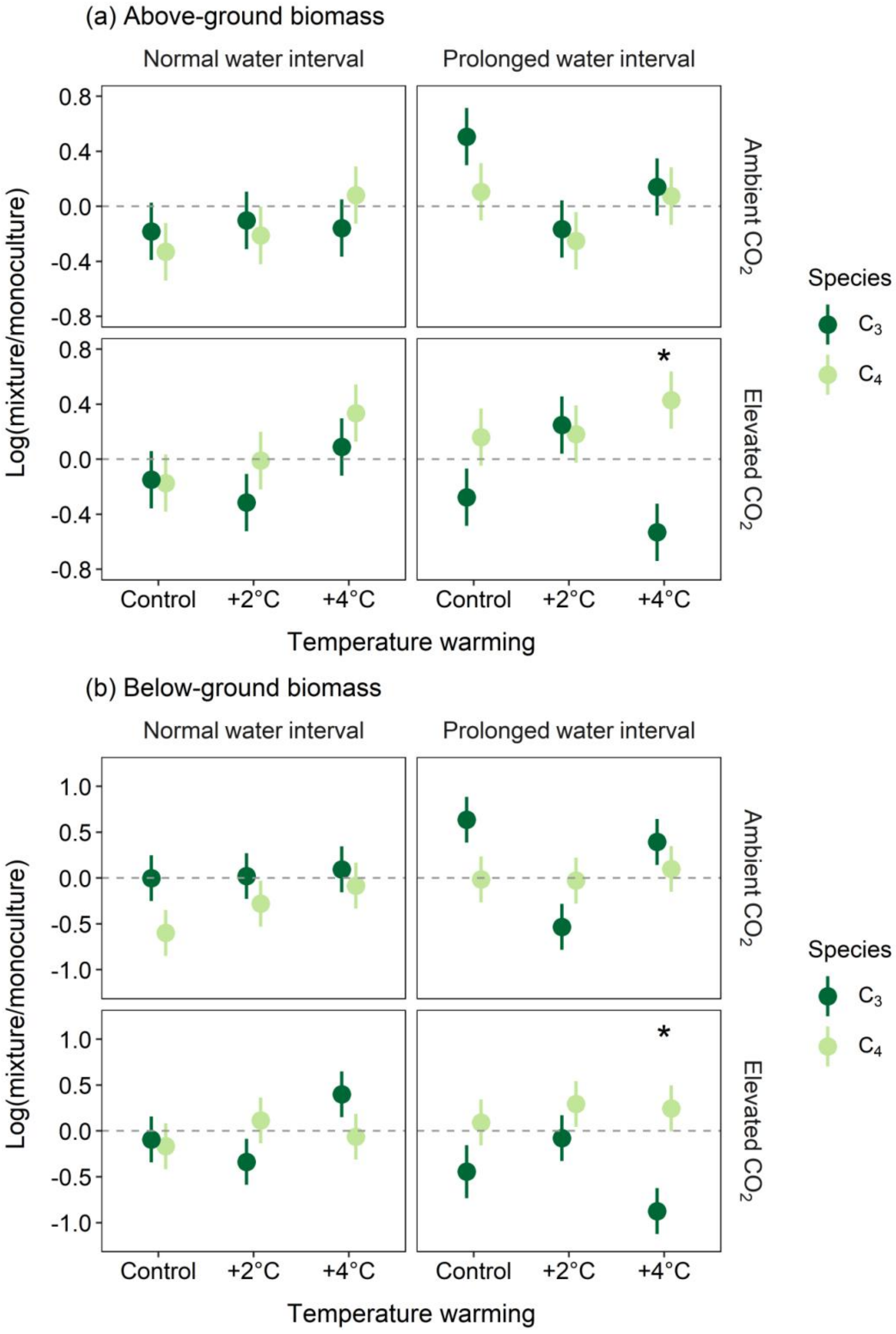
